# Genome sequence data reveal at least two distinct incursions of the tropical race 4 (TR4) variant of *Fusarium* wilt into South America

**DOI:** 10.1101/2022.01.17.476647

**Authors:** Paula H. Reyes-Herrera, Eliana Torres-Bedoya, Diana Lopez-Alvarez, Diana Burbano-David, Sandra L. Carmona, Daniel P. Bebber, David J. Studholme, Monica Betancourt, Mauricio Soto-Suarez

**Affiliations:** Corporación Colombiana de Investigación Agropecuaria - Agrosavia. C.I Tibaitatá. Km 14 vía Mosquera-Bogotá, Cundinamarca, Colombia; Biosciences, University of Exeter, Geoffrey Pope Building, Exeter, United Kingdom; Universidad Nacional de Colombia, Sede Palmira. Facultad de Ciencias Agropecuarias, Departamento de Ciencias Biológicas

## Abstract

The global banana industry is threatened by one of the most devastating diseases: Fusarium wilt (FWB). FWB is caused by the soil-borne fungus *Fusarium oxysporum* f. sp. *cubense* (*Foc*), which almost annihilated the banana production in the late 1950s. A new strain of *Foc*, known as tropical race 4 (TR4), attacks a wide range of banana varieties including Cavendish clones which are the source of 99% of banana exports. In 2019, *Foc* TR4 was reported in Colombia, and more recently (2021) in Peru. In this study, we sequenced three fungal isolates identified as *Foc* TR4 from La Guajira (Colombia) and compared them against 19 whole-genome sequences of *Foc* TR4 publicly available, including four genome sequences recently released from Peru. To understand the genetic relatedness of the Colombian *Foc* TR4 isolates and those from Peru, we conducted a phylogenetic analysis based on a genome-wide set of single nucleotide polymorphisms (SNPs). Additionally, we compared the genomes of the 22 available *Foc* TR4 isolates looking for the presence-absence of gene polymorphisms and genomic regions. Our results reveal that (i) the Colombian and Peruvian isolates are genetically distant, which could be better explained by independent incursions of the pathogen to the continent, and (ii) there is a high correspondence between the genetic relatedness and geographic origin of *Foc* TR4. The profile of present/absent genes and the distribution of missing genomic regions showed a high correspondence to the clades recovered in the phylogenetic analysis, supporting the results obtained by SNP-based phylogeny.

## Introduction

The global banana export industry generates around 14.5 billion USD per year (FAOSTAT 2020). Approximately 51% of global production is of the Cavendish cultivar (Lescot 2018). Latin America and the Caribbean (LAC) constitutes the world’s most important exporting region for bananas. During 2018, the total production volume of bananas in LAC was estimated at 30 million tonnes and 13 million tonnes were exported to developed countries, mainly the United States of America and the European Union. In Colombia, banana is the country’s third most important export crop after coffee and flowers. Approximately 91% of the national banana production is destined for export. Five departments concentrate almost 75% of the national production. In 2019, Antioquia was the department with the largest production of banana, with a cultivated area of 40,000 ha (42% of the total cultivated area of the nation), followed by Magdalena (14%), Nariño (9%), Valle del Cauca (7%) and La Guajira (3%) (Statistics home 2014).

Fusarium wilt of banana (FWB), one of the most devastating diseases of bananas, is threatening the global export industry. The disease, caused by the soil-borne fungus *Fusarium oxysporum* f. sp. *cubense (Foc)*, was first reported in Australia in 1876 (Bancroft 1876). The devastation caused by F*oc* race 1 (R1) was mitigated in the late 1950s by the substituting Gros Michel with resistant Cavendish cultivars, which now account for nearly all banana exports. However, a newly emerging lineage of *Foc*, known as tropical race 4 (TR4) corresponding to vegetative compatibility group VCG 01213/16, attacks Cavendish clones and a wide range of other banana varieties. *Foc* TR4 was first detected in Taiwan in 1967 (Hwang and Ko 2004; Su et al. 1977), after most likely being introduced on infected plants from Sumatra, Indonesia, and subsequently spread widely in banana-producing countries (Ploetz et al. 2015; Ploetz 2015). *Foc* TR4 was restricted to Australasia from 1990 when it was formally recognized until 2013 when it was first reported in Jordan, and Mozambique (Butler 2013). In 2015 it emerged in Lebanon, Oman, India and Pakistan (Viljoen et al. 2020; Ordoñez et al. 2016; Thangavelu et al. 2019; Ploetz et al. 2015). Between 2017 and 2019, *Foc* TR4 was found in Laos, Vietnam, Myanmar and Thailand (Ordoñez et al. 2016; García-Bastidas et al. 2014; Acuña et al. 2021; Chittarath et al. 2018; Hung et al. 2018; Zheng et al. 2018; Latest Pest Reports 2019). According to official information, *Foc* TR4 is currently confirmed in 22 countries (CABI ISC 2021), predominantly in South and Southeast Asia. In 2019, the pathogen was reported for the first time in Colombia, reaching out to the American continent (García-Bastidas et al. 2020; Viljoen et al. 2020), and more recently in Peru (Acuña et al. 2021). *Foc* TR4 was detected on a banana plantation in the northeastern region of La Guajira, Colombia. Currently, eleven farms in La Guajira and one in Magdalena are confirmed for the presence of *Foc* TR4. Consequently the 0.32 % of banana producing region in the country is under quarantine due to the presence *Foc* TR4.

Understanding the phylogenetic relationships among geographically disparate isolates can provide clues about a pathogen’s chains of transmission. An understanding of the genetic diversity and relationships with other organisms is important for rational design of molecular assays for detection and identification. Taxonomy is important for implementation of control measures such as notification and quarantine.

Recently, the use of high-throughput genome-sequencing technologies has made important contributions to studies on *Foc* TR4, particularly on genetic diversity (Maymon et al. 2020; Ordonez et al. 2015) and phylogeographical analysis (Zheng et al. 2018). Ordoñez and colleagues (Ordoñez 2018; Ordonez et al. 2015) performed a hierarchical clustering analysis based on 4,298 DArTseq markers showing a limited genetic diversity between multiple *Foc* TR4 isolates from countries in the Middle East, Asia and Oceania (China, Indonesia, Jordan, Lebanon, Malaysia, Pakistan, Philippines and Australia). Genomic comparison of *Foc* TR4 isolates from the Greater Mekong subregion (Zheng et al. 2018) identified three geographically distinct clusters, one of which constituted isolates from Laos, Vietnam, Myanmar and China; this suggested that the source of infection in the Greater Mekong subregion probably originated from China. Most recently, a phylogeographical study conducted by Maymon and colleagues (Maymon et al. 2020), using SNPs across the whole genome, principal component analyses (PCA) and hierarchical clustering, claimed a strong similarity between the Colombian isolates and the Indonesian isolate II-5. The authors argued that this suggests that the pathogen most likely spread to Colombia from Indonesia (Maymon et al. 2020). A recent genomic comparison of Indian *Foc* isolates belonging to races 1, 2 and 4 revealed differences in the repertoire of *SIX* genes in Indian *Foc* TR4 compared with *Foc* TR4 isolated elsewhere (Raman et al. 2021).

Previous work (Maymon et al. 2020) suggested that the source of the Colombian *Foc* TR4 outbreak might be Indonesia. It is likely that the *Foc* TR4 lineage first arose in that part of the world (Bentley et al. 1998; Vézina 2014) and in that sense, Indonesia might ultimately be the origin of all *Foc* TR4. However, it remains unclear as to what routes *Foc* TR4 has taken to disseminate across and between continents. Furthermore, misattribution of the pathogen’s source may have important economic and political consequences; so, any such claims require careful scrutiny. Therefore, the aims of this study were (i) to evaluate the SNP genetic diversity between Colombian isolates and those from elsewhere, including genome sequences for three additional Colombian isolates, four from Peru and other sequence data that were not available at the time of the previous study (Maymon et al. 2020), and (ii) to identify new genomic differences/similarities between TR4 isolates able to support the SNP-based phylogeny. For this, we sequenced the genomes of the three additional *Foc* TR4 isolates using Oxford Nanopore Technologies’ MinION platform. Our genomic comparison analyses revealed that outbreaks in Peru and Colombia are genetically distinct and likely have different origins. We generate a catalogue of genes/regions whose presence is variable among *Foc* TR4 pathogen individuals that will be useful resource for future study.

## Results and Discussion

The *Foc* TR4 variant of Fusarium wilt has been detected in two South American countries, namely Peru and Colombia. This lineage of the pathogen likely emerged initially in Indonesia or Malaysia (Bentley et al. 1998; Vézina 2014) and has subsequently undergone a number of intercontinental transmission events. Each of these events presumably involved a founder population that represents a tiny sample of the pathogen population’s genetic diversity, leading to a genetic bottleneck at each introduction to a new geographical location. Until recently, *Foc* TR4 was unknown in Latin America. This raises the question of whether recent outbreaks in the two South American countries are linked in a direct chain of transmission or whether they represent separate introductions. The ‘bottleneck’ effect of introduction from the *Foc* TR4 population in its centre of origin predicts that directly linked outbreaks would involve genetically similar pathogen individuals, whereas independent samples from the that original population would be relatively divergent from each other. We compared isolates from Colombia with previously sequenced isolates from elsewhere, using genome-wide sequence data to maximise the resolution of the genetic relationships.

### Genome sequencing of Colombian *Foc* TR4 isolates

We generated approximately 17 Gb of long-read data for each Colombian *Foc* TR4 isolate. The sequence reads have been deposited in the Sequence Read Archive under the BioProject accession number PRJNA774343 (BioSample accession numbers SAMN22562322, SAMN22562323 and SAMN22562324). These genomic reads were aligned against the UK0001 *Foc* TR4 reference genome sequence and the resulting alignments were used for calling SNPs for phylogenetic analysis and for surveying differentially present/absent genes.

### Presence/absence of *SIX* genes in the Colombian *Foc* TR4 genomes

During the last ten years, several studies have developed molecular markers for detection of *Foc* TR4 (Matthews et al. 2020; Magdama et al. 2019; Ndayihanzamaso et al. 2020; Dita et al. 2010; Aguayo et al. 2017; Lin et al. 2013; Li et al. 2013; Carvalhais et al. 2019). This raises an important question: Are the existing molecular markers present in the Colombian *Foc* TR4 genomes? We investigated whether the secreted in xylem *SIX* genes (*SIX1* – *SIX13*) were present/absent in Colombian *Foc* TR4 isolates. For this, a BLAST analysis was carried out using *SIX* gene sequences against the 190098, 190203 and 03242 Colombian genome assemblies. All sequences were present in all three Colombian *Foc* TR4 genomes. *SIX-gene* homologues from *SIX1* to *SIX13* showed different levels of similarity, ranging from 98 to 100%, to those of our sequenced isolates (Supplementary Table S1). Interestingly, *SIX6* and *SIX9* genes presented low percentage similarities of 91 and 88%, respectively.

### Colombian isolates constitute a distinct clade distinct from Peruvian isolates

We identified 671 single-nucleotide sites in the *Foc* TR4 genome that showed variation and for which the allele could be unambiguously determined in every one of the examined genomes. On the basis of these 671 SNPs (Supplementary File S2), we constructed the maximum-likelihood phylogeny shown in Figure 1. This phylogenetic tree displayed a clear correspondence between genetic relatedness and geographic origin. For example, there is a clade comprising isolates from the Middle East. Notably, isolates from South America are distributed among two distinct clades. The six isolates from Colombia comprise a single clade; similarly, the four isolates from Peru comprise another distinct clade. Colombian isolates are genetically much more distant from Peruvian isolates than they are from isolates collected in the Middle East and the United Kingdom. This phylogeographic pattern is not consistent with a single introduction of *Foc* TR4 from its centre of origin into South America and subsequent spread within the continent. Rather, it is better explained by separate, independent incursions into Colombia and Peru.

**Figure 1.**
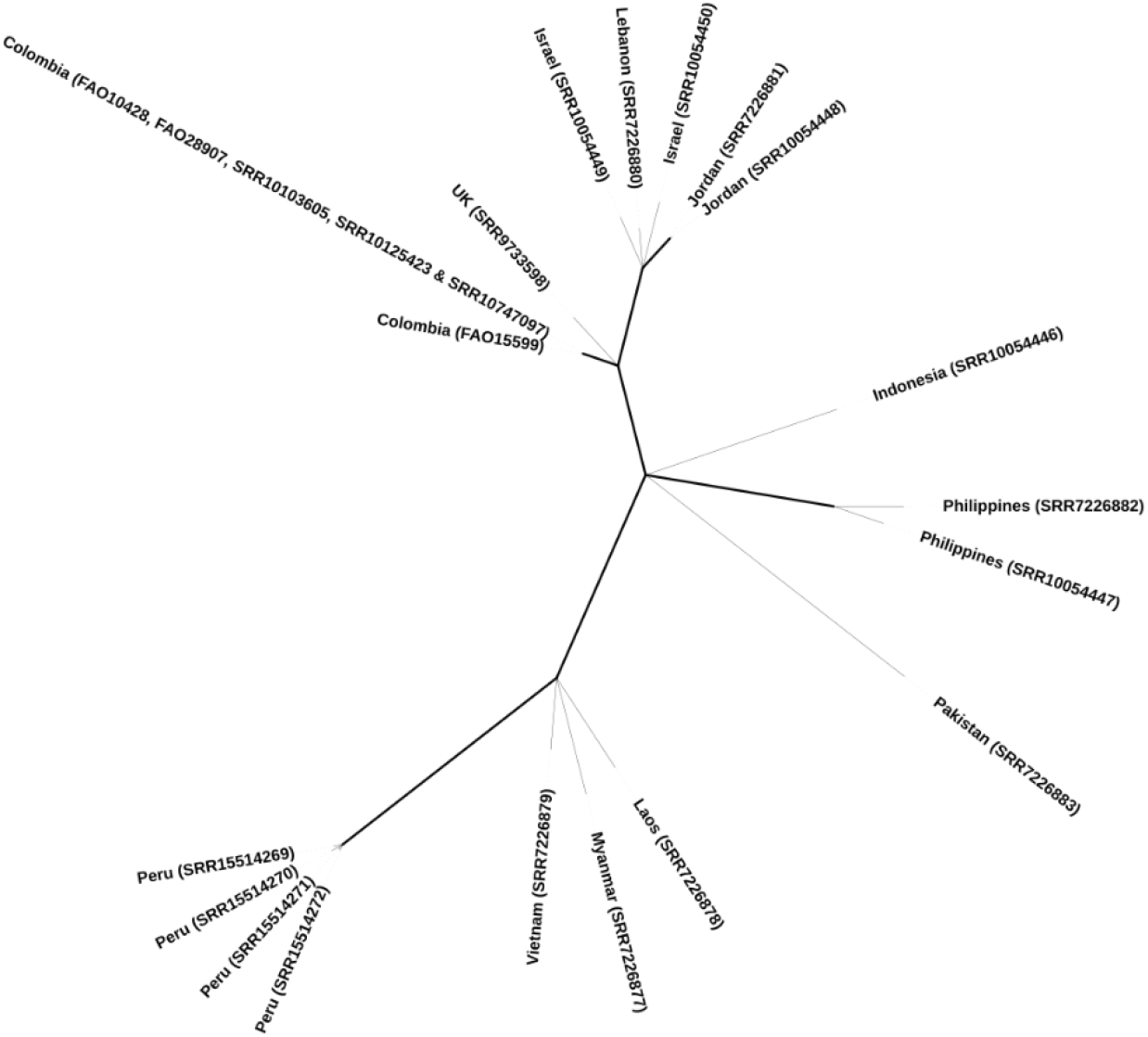
Maximum-likelihood phylogeny of genome-sequenced Foc TR4 isolates from diverse geographical sources. The tree is based on 671 single-nucleotide polymorphisms and built using IQ-Tree (Nguyen et al. 2015). We obtained branch supports with the ultrafast bootstrap (Hoang et al. 2018).

Our phylogenetic analysis revealed that the six Colombian isolates are much more closely related to each other than they are to any previously sequenced isolates from other geographic locations. In other words, there is greater genetic differentiation between than within geographical regions. Furthermore, there is no close relationship between Colombian isolates and isolates from elsewhere; Colombian isolates are genetically approximately equidistant to each of the other isolates. Therefore, there is no genetic evidence of a direct source of the incursion of *Foc* TR4 into the Americas; the lineage comprising the Colombian isolates probably diverged from the lineages isolated elsewhere prior to the emergence of *Foc* TR4 outside of its centre of origin (whose location is unknown but likely to lie in Indonesia and/or Malaysia). The phylogenetic tree is consistent with several independent intercontinental transmissions of *Foc* TR4 from its origin. Similarly, within China and SE Asia, all sequenced isolates are closer to each other than to isolates from elsewhere, suggesting a single egress of *Foc* TR4 into that region. Similarly, most of the isolates from countries in the Middle East are genetically close and may have arisen from a single inoculum. In conclusion, the phylogenetic analysis is consistent with a single source of *Foc* TR4 into the Americas and shows no evidence that this is derived from the *Foc* TR4 populations seen in other regions where it has emerged.

### Patterns of gene content are broadly consistent with phylogeny

Based on alignments of genomic reads from *Foc* TR4 isolates against the UK0001 reference genome sequence, we identified 615 gene presence-absence polymorphisms. These are tabulated as an Excel spreadsheet in Supplementary File S3. The distributions of these variable genes across the sequenced isolates were broadly consistent with phylogeny. Clustering of the genomes based on their profile of present/absent genes showed clusters that corresponded to the clades recovered in the phylogenetic analysis (Figure 2), revealing dozens of genes whose presence or absence is characteristic of specific *Foc* TR4 clades. This opens the future possibility of developing molecular typing assays to assign isolates to lineages without the need for whole-genome sequencing. Furthermore, it is notable that several genes show presence-absence polymorphism within the Colombian clade, suggesting that gene deletions have taken place very recently during the epidemic, some of which might be adaptive as the fungus finds itself in a new environment with new host genotypes. The biological significance of these differentially present genes is an avenue for future investigation.

**Figure 2.**
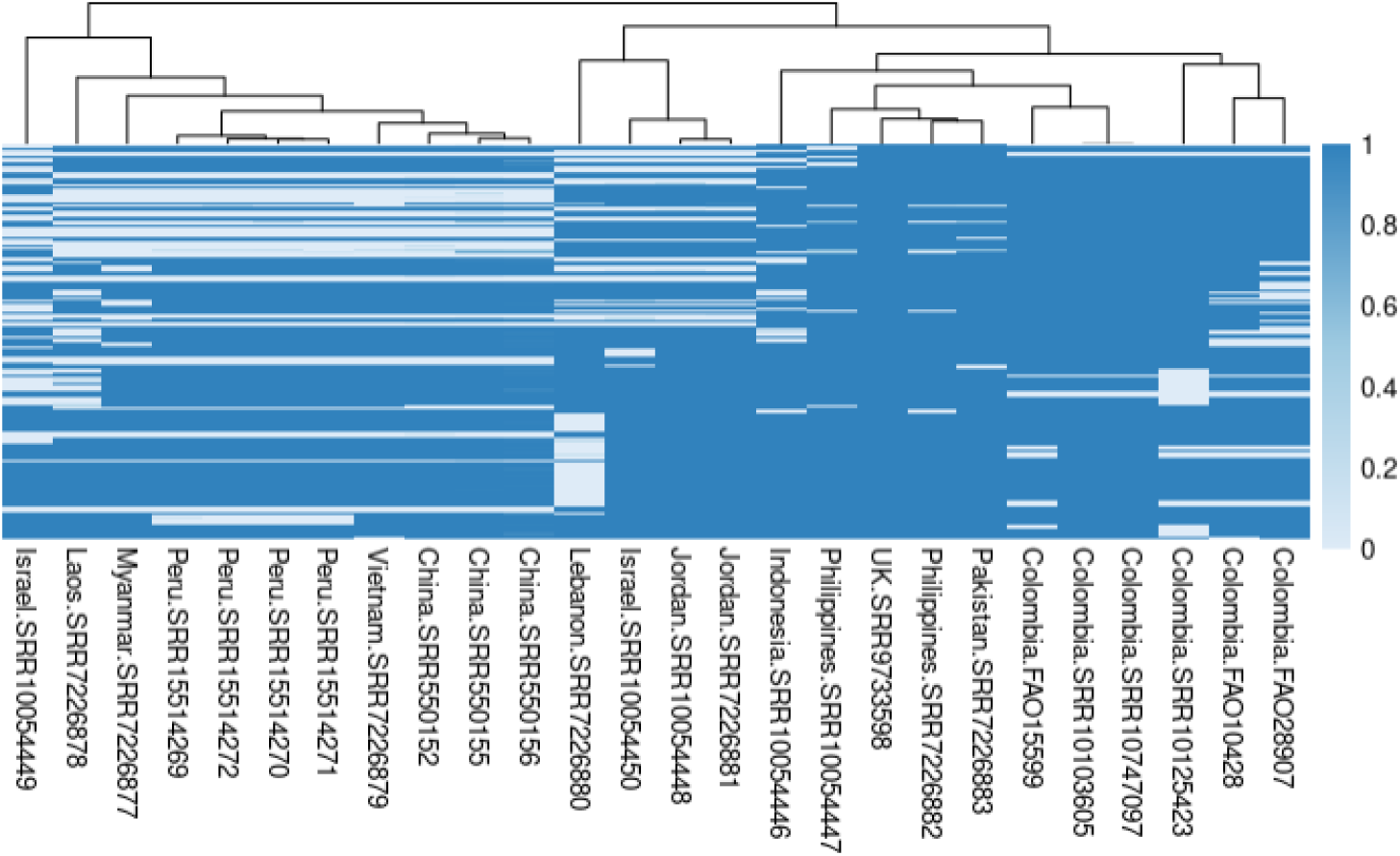
Heatmap showing a comparison of gene content of Foc TR4 genomes. The rows represent 615 predicted protein-coding genes in the UK0001 reference genome that are absent from at least one TR4 genome; that is, differentially present/absent genes. The presence or absence of each gene was assessed in each genome based on breadth of coverage by genomic sequencing reads, using the coverageBed tool (Quinlan and Hall 2010); this generates a value between zero (no reads, i.e. absent) and one (completely covered by reads, i.e. present). The columns (i.e. genomes) are ordered according to by complete linkage clustering. Rows (i.e. genes) are ordered according to genomic location.

### Comparative genomic analysis reveals sequence gaps shared by *Foc* TR4 isolates from specific geographic locations

A comprehensive genomic analysis can identify components of the *Foc* TR4 genome that might complement the information provided by the SNP-based phylogeny. In this study, the whole genome assemblies were compared to the published genomes of *Foc* TR4 (Supplementary Table S4). We compared the genomes of 22 publicly available *Foc* TR4 isolates with the reference genome assembly of UK0001 (Figure S1). The comparison between the *de novo* assembled Colombian *Foc* TR4 shows that they do not differ from each other. However, when comparing all *Foc* TR4 genomes, we observed several major gapped regions located on contigs VMNF0100005.1, VMNF0100007.1, VMNF01000013.1 and VMNF01000014.1 (Figure 3). Specifically, genomic regions that are present on the UK0001 reference genome are missing in (i) the Colombian *Foc* TR4 isolates (contigs VMNF0100005.1 andVMNF0100007.1); (ii) Middle Eastern isolates from Jordan, Israel and Lebanon (contigs VMNF01000013.1 and VMNF01000014.1); (iii) isolates from China, Vietnam, Myanmar, Laos and Peru (VMNF01000014.1). Thus, this comparative genomic analysis also supports the SNP-based phylogeny.

**Figure 3.**
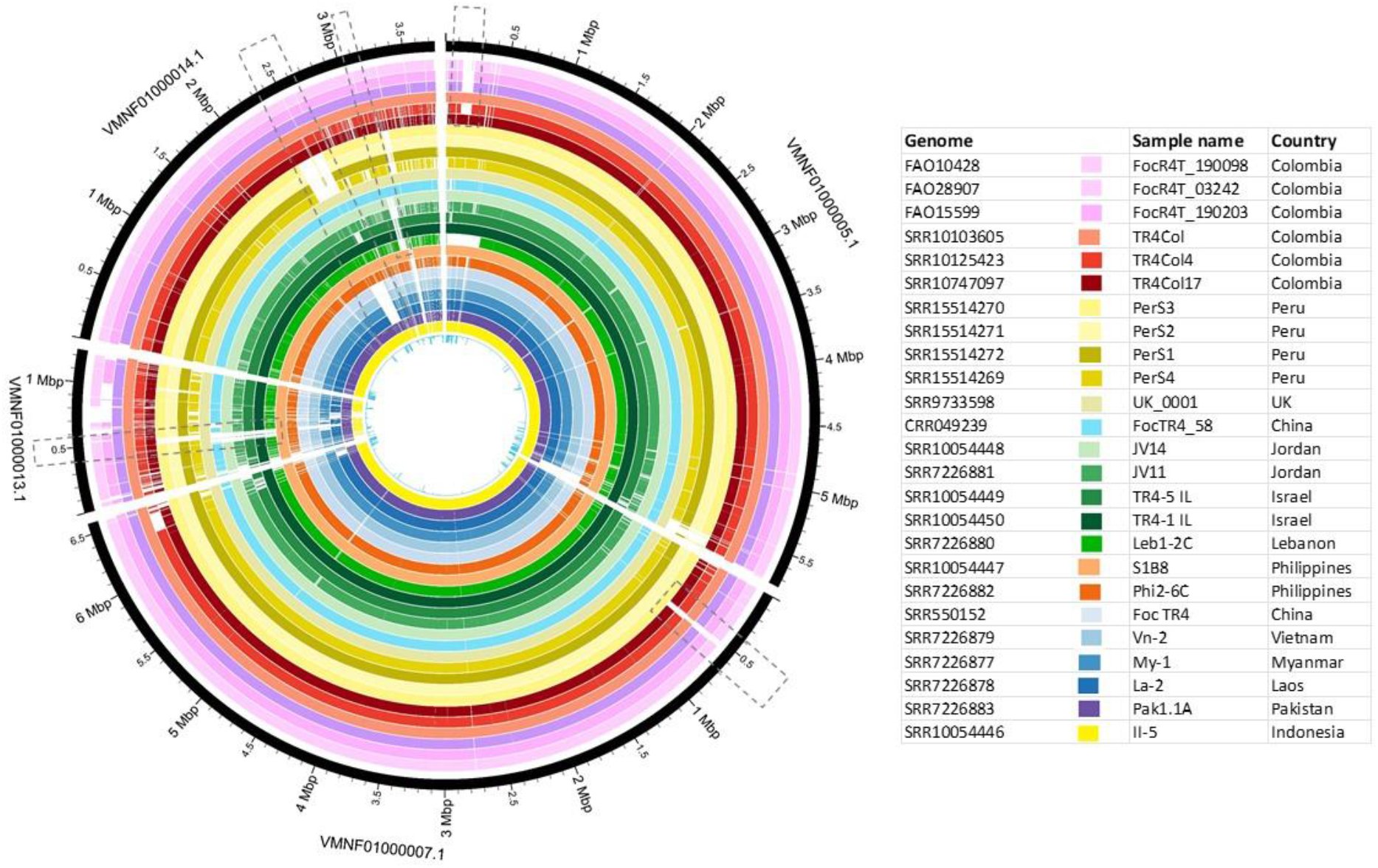
Comparative genomic analysis of 22 *Foc* TR4 genomes. Circle map showing contigs VMNF0100005.1, VMNF0100007.1, VMNF01000013.1 and VMNF01000014.1. The dotted rectangles indicate the missing genomic regions.

### Conclusion

Phylogeographic relationships among *Foc* TR4 worldwide isolates are required to infer the routes of TR4’s transmission between and within continents. In this work, we sequence and obtained a nearly complete genome assembly of three isolated of *Foc* TR4, obtained from banana plantations of la Guajira in Colombia. Our analysis suggests that Colombian isolates are more closely related to those from the UK and the Middle East and less to those from Perú. However, the divergences between those three lineages likely occurred prior to the global emergence of *Foc* TR4 outside its centre of origin, based on the genetic distances seen within geographically distinct lineages. Comparative genomic analysis revealed missing genomic regions that were shared between *Foc* TR4 isolates belonging to the same clade, also confirming a correspondence between genetic relatedness and geographic origin.

## Materials and methods

### Genome sequence data from public repositories

We used the previously published UK0001 genome (Warmington et al. 2019) as the reference sequence for alignment-based analyses as that genome sequence was assembled using long reads and is therefore of high quality. We used genome sequencing reads for previously reported *Foc* TR4 genome sequencing projects from the Sequence Read Archive (SRA) accession numbers SRR10103605, SRR10125423, SRR10747097, SRR9733598, SRR7226880, SRR10054450, SRR10054449, SRR7226881, SRR10054448, SRR10054446, SRR10054447, SRR7226882, SRR7226883, SRR7226879, SRR15514269, SRR15514270, SRR15514271, SRR15514272, SRR550155, SRR550152, SRR7226878 and SRR7226877 (Guo et al. 2014; Leinonen et al. 2011; Kodama et al. 2012; Warmington et al. 2019; Maymon et al. 2020; Acuña et al. 2021; Zheng et al. 2018).

### Colombian *Foc* TR4 isolates

The *Fusarium oxysporum* f. sp. *cubense* TR4 isolates 190098, 03242 and 190203, isolated from symptomatic Cavendish banana from Dibulla, La Guajira state (Colombia) were provided by Instituto Agropecuario Colombiano (ICA). The isolates were maintained in potato dextrose agar (PDA) at 27 °C.

### Extraction of genomic DNA

For high molecular weight (HMW) DNA extraction, the *Foc* TR4 isolates were grown on PDA plates for seven days at 28°C. Then, propagules were harvested and transferred to Czapek dox medium and incubated for six additional days to produce fungal mycelia. Fungal mycelium was obtained by filtering through two layers of Miracloth and washed twice with 10 mL sterile distilled water. Then, fungal mycelium was freeze-dried overnight and ground in a mortar with a pestle. Five hundred milligrams of ground mycelium were incubated for one hour at 65°C with 800 μL fresh DNA extraction buffer and 15 μl de RNAse (10mg/μl). DNA extraction buffer was prepared by mixing 2.5 volumes of solution A (0.35 M Sorbitol, 0.1 M Tris-base, 5 mM EDTA pH 7.5), 2.5 volumes of solution B (0.2 M Tris, 0.05 M EDTA, 2 M NaCl, 2% CTAB), 1 volume of Sarkosyl (10% w/v) and 1% β-mercaptoethanol. To separate the organic phase, 400 μL of phenol/chloroform/isoamyl alcohol (25:24:1) was added, vortexed for 5 minutes and incubated at room temperature (RT) for 5 minutes before centrifugation at 16,000 g for 15 min. Two chloroform extractions were used on the aqueous phase by adding 0.5 volumes and centrifuge at 16,000 g for 5 min at RT each time. The aqueous phase was mixed with 10 volumes of 100% ice-cold ethanol, incubated for 30 min at RT, and the precipitated DNA was collected in a new tube using a disposable inoculation loop. Collected HMW DNA was washed twice with 1 mL 70% ice-cold ethanol and the air-dried DNA was resuspended in nuclease-free water and conserved at 4°C. The DNA quality, size and quantity were assessed by spectrometry in a NanoDrop 2000 Spectrophotometer (Thermo Fisher Scientific, Wilmington, USA), electrophoresis in agarose gel and fluorometry in a Qubit Fluorometer v2.0 (Life Technologies, Thermo Fisher Scientific Inc.)

### DNA sequencing

Sequencing library was prepared with the Ligation Sequencing Kit (SQK-LSK109) according to the manufacturer instructions (Oxford Nanopore Technologies, Oxford, UK) using 22 ng HMW DNA. An R9.4.1 flow cell (Oxford Nanopore Technologies, Oxford, UK) was loaded and run for 48 hours. Base calling was performed using Guppy from MinKNOW (version 4.0.21; Oxford Nanopore Technologies).

### Genome assembly

For the *de novo* assembly of Oxford Nanopore Technologies data from three newly sequenced Colombian isolates, we used Canu 2.0 (Koren et al. 2017) and Flye 2.8.2 (Kolmogorov et al. 2019) followed by several iterations of Racon v1.4.3 (Vaser et al. 2017) and medaka 1.2.1 (Oxford Nanopore Technologies Ltd. 2018) to obtain a consensus sequence. The best assembly for each isolate was selected according to most of the metrics such as BUSCO score (Simão et al. 2015), QUAST genome statistics (Gurevich et al. 2013) and Qualimap (García-Alcalde et al. 2012). In addition, BlobTools2 (Challis et al. 2020) as used to integrate to include coverage, BUSCO, Blast, and DIAMOND v2.0.11 (Buchfink et al. 2015) and do contaminant screening and genome assessment.

For most of the previously sequenced genomes, assemblies were available in the public databases. However, for several isolates, only unassembled sequence reads were available. Assemblies of isolates SRR10125423, SRR10747097, SRR550152, SRR550155, SRR7226877, SRR7226878, SRR7226879, SRR7226880, SRR7226881, SRR7226882, and SRR7226883 were performed using SPAdes and evaluated with Qualimap (García-Alcalde et al. 2012). For genomes with low numbers of reads (SRR10054447, SRR10054448, SRR10054449, SRR10054450, SRR10103605, SRR15514270, SRR15514271, and SRR15514272) we aligned the reads to the reference genome using Bowtie2 (Langmead and Salzberg 2012) and used the mpileup tool in Samtools (Li et al. 2009) to obtain a consensus sequence from the alignment. MUMmer4 was used to align the whole genome assemblies to the reference genome UK0001. Then, we used Circos to visualize the alignments in a circular representation.

### Aligning sequence reads against reference genome sequence

To mitigate problems arising from incompleteness and errors in *de novo* assembly of short sequence reads, we used a genome-comparison strategy based on aligning sequencing reads against a high-quality reference genome sequence. For this assembly-free genomic comparison, we acquired short-read whole-genome Illumina sequence data as FastQ files (Cock et al. 2010) for *Fusarium oxysporum* f. sp. *cubense* from the SRA database (Kodama et al. 2012). The quality of the sequencing data was evaluated using FASTQC (Andrews n.d.). Reads with low quality or containing adaptor sequences were trimmed using Trim Galore (Babraham Bioinformatics - Trim Galore! 2022) or Canu (Koren et al. 2017) as appropriate to the sequencing method that generated the data. The sequences were aligned against the reference genome of isolate UK0001 (GenBank:GCA_007994515) using the Burrows Wheeler Aligner (BWA) (Li and Durbin 2010, 2009) and Minimap2 (Li 2018). The alignments were evaluated with Qualimap (García-Alcalde et al. 2012).

### SNP-calling and phylogeny reconstruction

We used a genome-wide survey of SNPs towards understanding the relationship between Colombian *Foc* TR4 isolates and those from other geographical locations.

Single-nucleotide sites that showed sequence variability between *Foc* TR4 isolates (i.e. candidate SNPs) were identified using Pilon (Walker et al. 2014). There was some variation in the level of confidence in the nucleotide sequences at these candidate SNPs. For example, at some sites, there was not a consensus among the multiple sequence reads aligned at that site. Therefore, candidate SNPs were filtered to retain only high-confidence SNPs with read-consensus above 95%, using a Perl script available at https://github.com/davidjstudholme/SNPsFromPileups. Full details of the command lines are given in Appendix 1 at the end of this document. Genomes were assigned to genetic types (haplotypes) according to the combination of nucleotide-states found at each of the high-confidence SNPs. These were represented as files in FastA and Nexus formats. A PhyML tree was generated in IQ-TREE (Nguyen et al. 2015) using the GTR model (Tavare 1986; Gouy et al. 2010; Guindon et al. 2010). The robustness of the phylogeny was assessed using 1000 bootstrap replicates.

### Comparison of gene content

We identified genes in the UK0001 reference genome that were absent from one or more sequenced isolates (i.e. presence-absence polymorphisms) from the BWA alignments using the *coverageBed* tool in BEDtools (Quinlan and Hall 2010). This tool reports the breadth of coverage by aligned sequence reads for each gene. This approach, based on aligning reads against a reference genome, avoids problems arising from incompleteness of *de-novo* genome assemblies.

## Author contributions

The study was conceived by MS-S and MB. The experiments were supervised by MS-S and MB. DNA extraction, library preparation and sequencing were conducted by SLC and DB-D. The analyses performed in the study were conceived by MS-S, DL-A, ET-B, DJS and PHR-H. Data analyses were completed by PHR-H, DL-A and ET-B. DPB and DJS supervised ET-B’s bioinformatic analyses. PHR-H, DL-A, ET-B, MB, DPB, DJS and MS-S interpreted results. MS-S and DJS drafted a first version of this manuscript, edited by all other co-authors. All authors contributed to the article and approved the submitted version.

## Funding

This work was supported by the earmarked fund to Agrosavia from Colombian Ministry of Agriculture, grant number (1000734: An update on Musaceae family diseases).

The Colombian *Foc* TR4 isolates were registered in the National Collections Registry (RNC129) and was collected under the AGROSAVIA’s permit framework No.1466 from 2014, updated by the 04039 resolution on July 19th, 2018.

### Institutional Review Board Statement

Not applicable.

### Informed Consent Statement

Not applicable.

## Data Availability Statement

The data that supports the findings of this study are available in the supplementary information of this article. Any additional data will be available on request to the corresponding author. Genome sequence data have been deposited in the Sequence Read Archive and GenBank and are available via BioProject accession number PRJNA774343 (BioSamples SAMN22562322, SAMN22562323 and SAMN22562324) and PRJNA731180 (BioSamples SAMN19572426, SAMN19275239 and SAMN19275177).

## Supplementary materials

**Supplementary Figure 1.**
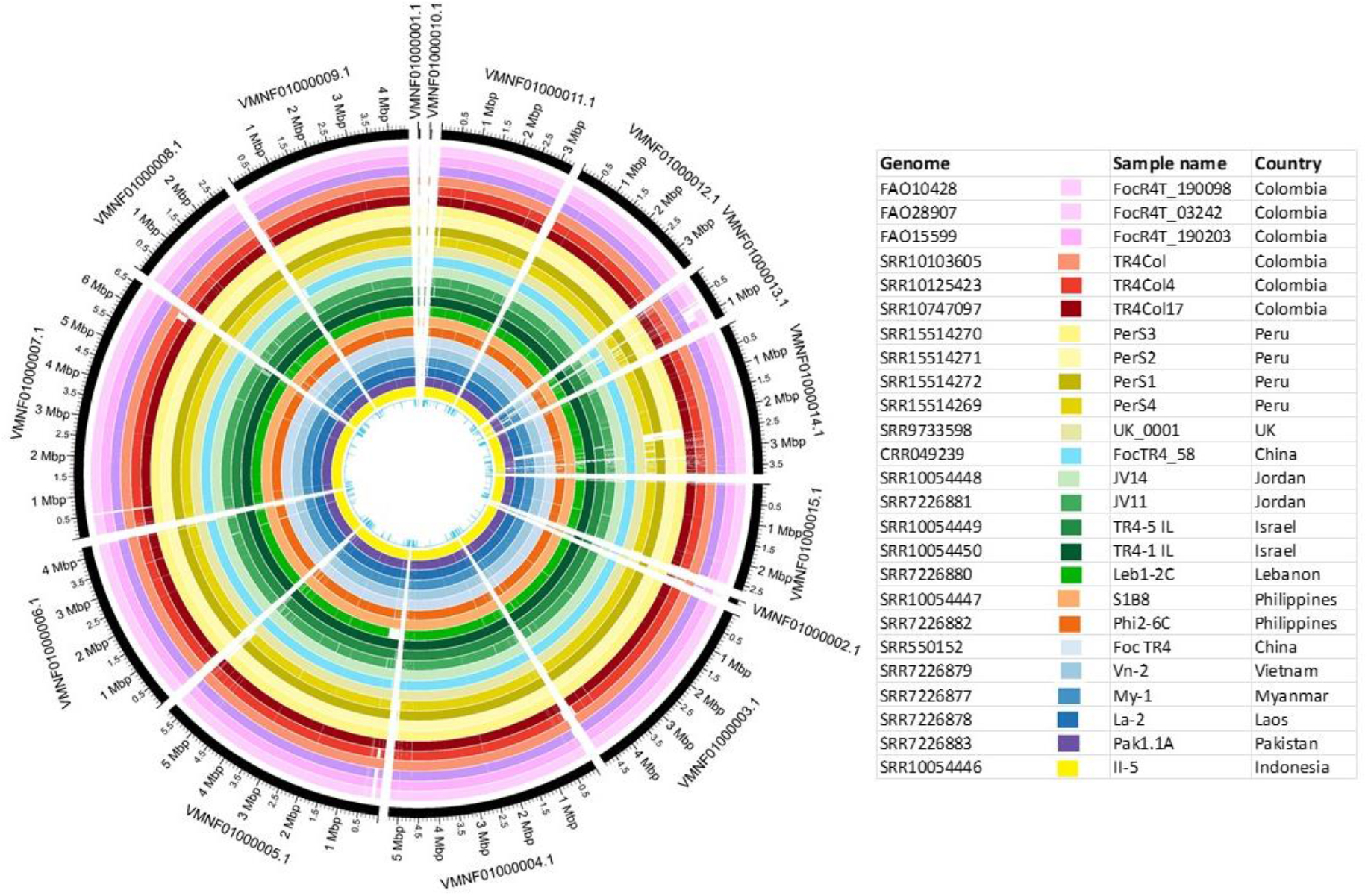
Circos plot for all the contigs alignment between the 22 *Foc* TR4 genomes and the Reference isolate UK0001 used as reference.

**Supplementary Table S1.** Summary of BLAST analysis using *SIX* genes against assembled *Foc* TR4 isolates of Colombian origin.

**Supplementary File S2.** Tabulated Excel spreadsheet containing the 671 SNPs 671 single-nucleotide sites in the *Foc* TR4 genome that showed variation.

**Supplementary File S3.** Tabulated Excel spreadsheet containing the 615 gene presence-absence polymorphisms identified after comparing genomic reads from *Foc* TR4 isolates against the UK0001 reference genome sequence.

**Supplementary Table S4.** Information on three newly assembled and 19 publicly available *Foc* TR4 genomes analysed in this study.

## Appendix 1: Command lines used for bioinformatics analysis

### Gene presence/absence polymorphisms, SNPs and phylogeny of TR4 genomes

#### Download the reference genome sequence

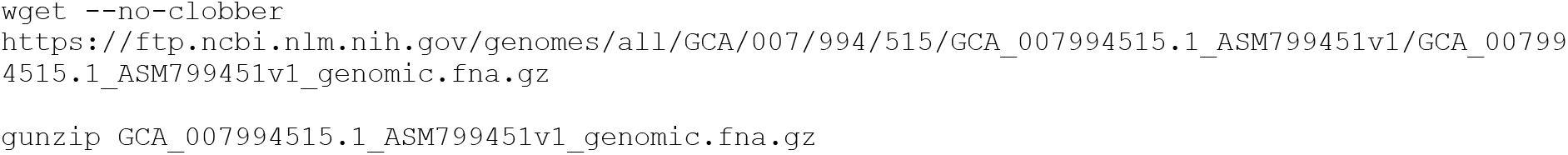

#### Generate the pileup files from BAM files

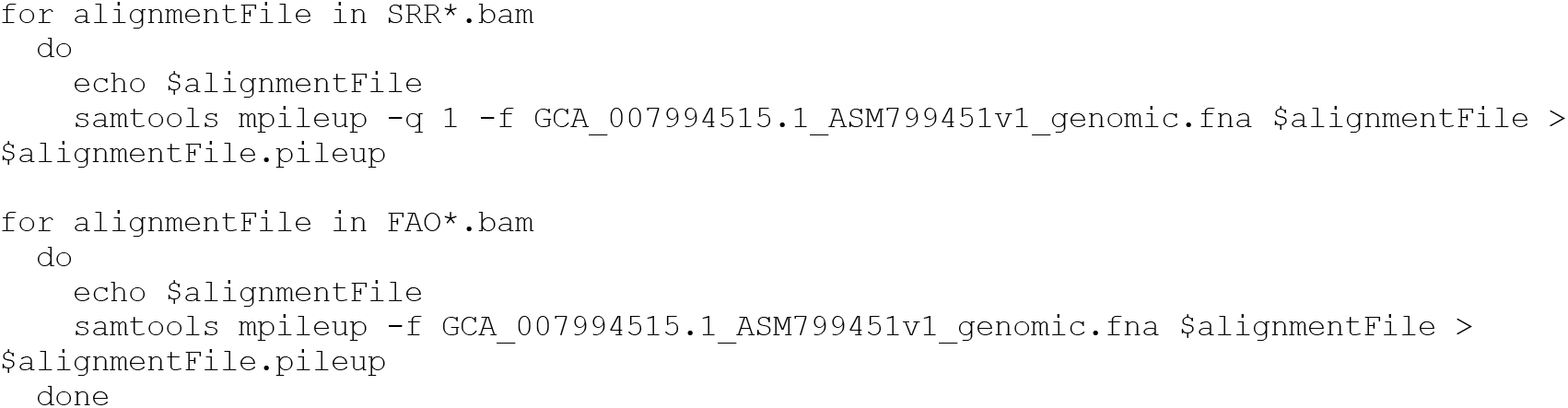

#### Index the BAM files

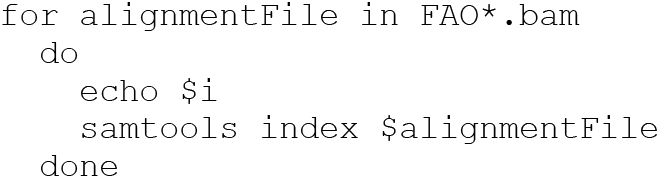

### Call SNPs

#### Identify candidate SNP sites using Pilon

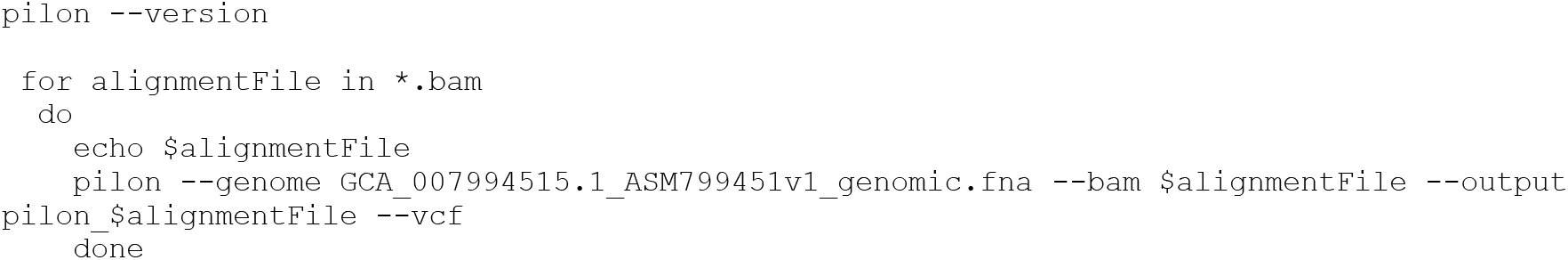

#### Filter the candidate SNPs

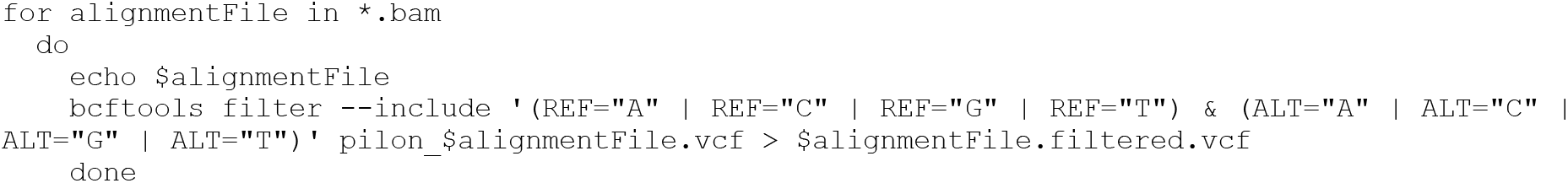

#### Get the SNP-calling scripts from GitHub

~~~
git clone https://github.com/davidjstudholme/SNPsFromPileups.git
~~~

#### Perform SNP-calling from pileup files

To minimise memory usage, we only consider candidate sites previously identified using Pilon.

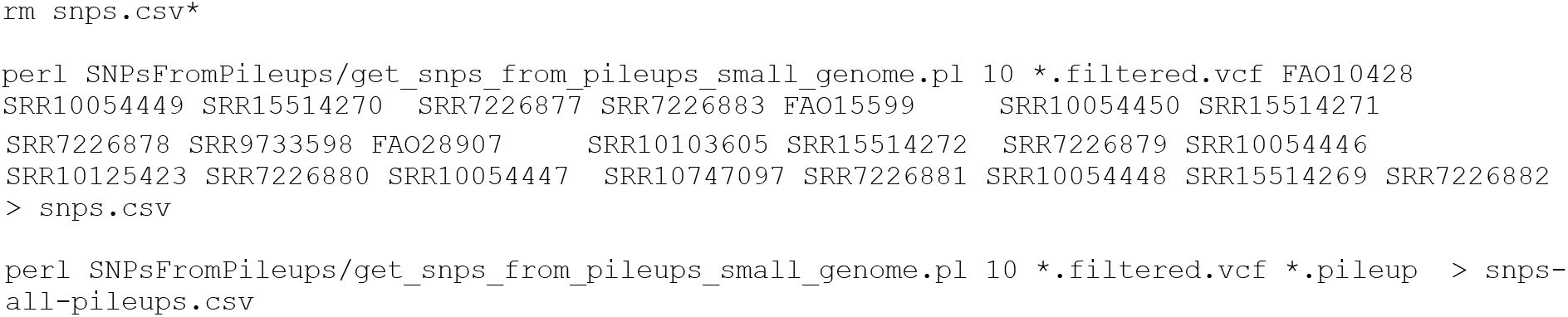

### Convert the SNPs into Nexus format for input into IQ-Tree

~~~
perl SNPsFromPileups/get_haplotypes_and_aligned_fasta_from_csv.pl snps.csv
~~~

### Perform phylogenetic analysis using IQ-Tree

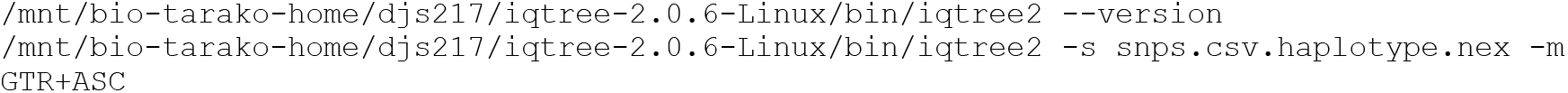

### Perform bootstrapping

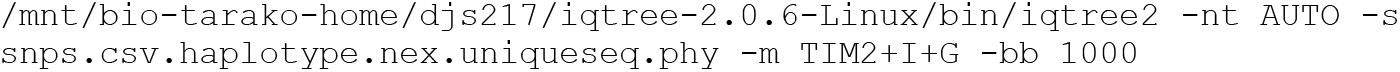

### Examine gene content using coverageBed from Bedtools

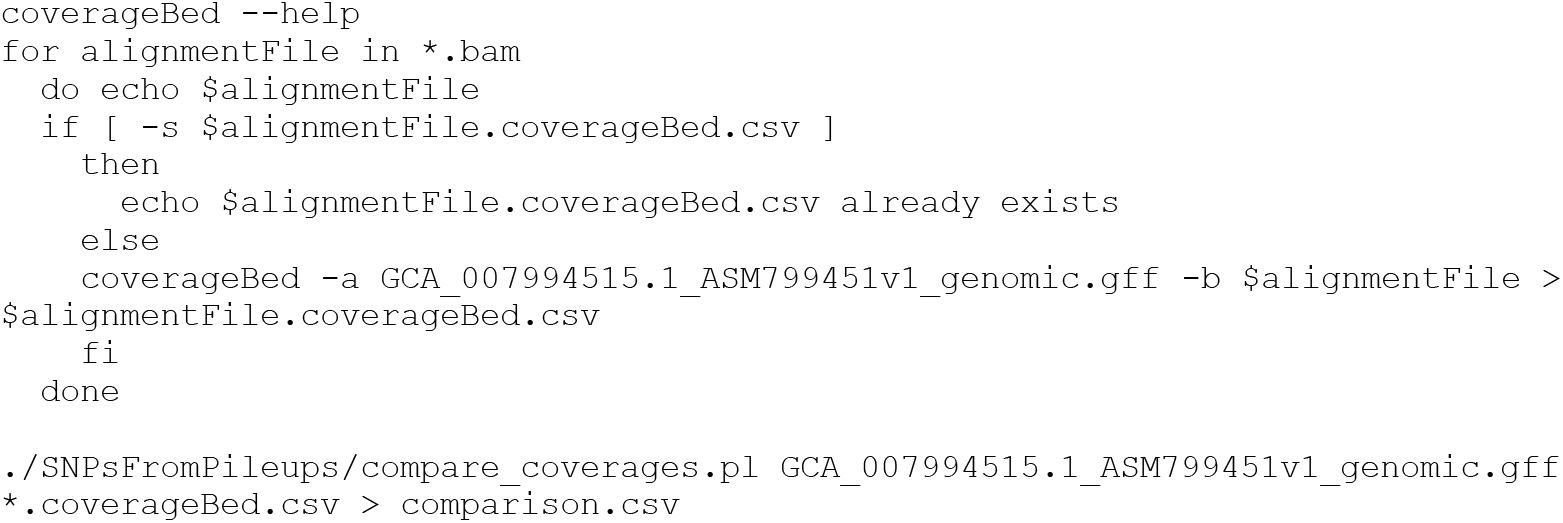

### Plot the variable genes as a heatmap

#### Install the packages

~~~
install.packages(‘pheatmap’)
~~~

#### Load the packages

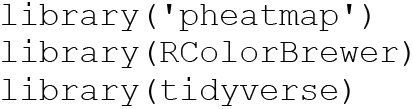

#### Make the plot

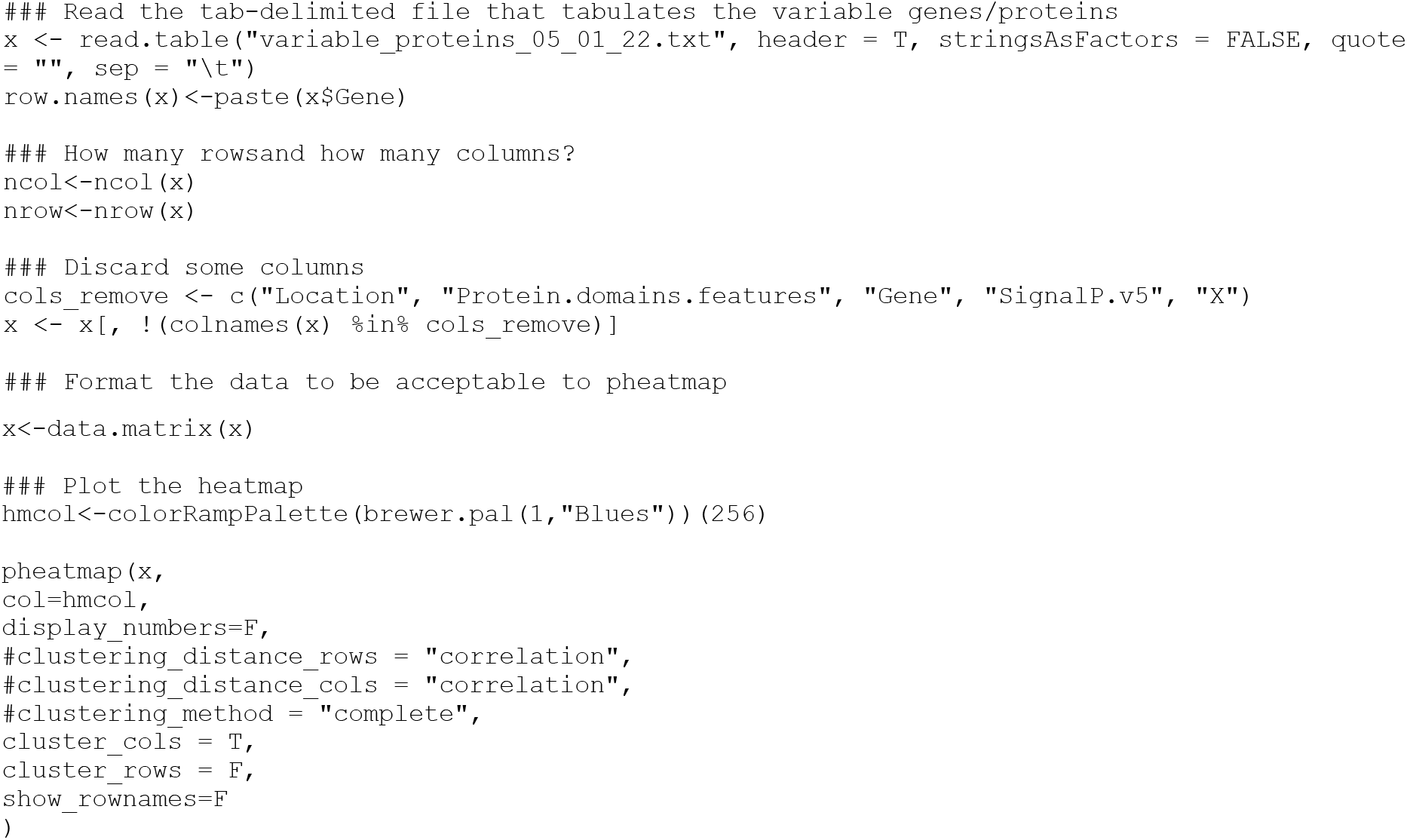

